## Supplementary Information for "Independent control of mean and noise by convolution of gene expression distributions"

### Supplementary Figures

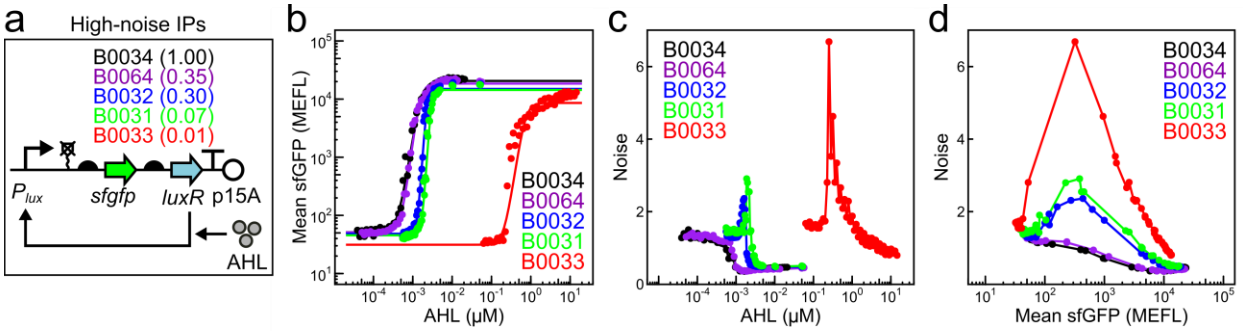

**Supplementary Figure 1**. **Modulating output noise of LuxR-based positive feedback IPs with RBS strength**. (**a**) LuxR based positive autoregulation IPs with different RBSs (including B0033 not shown in the main text) used for generation of high noise. AHL-mean (**b**) AHL-noise (**c**) and mean-noise (**d**) transfer functions of IPs from (**a**). The B0033 variant generated purely bimodal distributions and was therefore not used as $IP_h$.

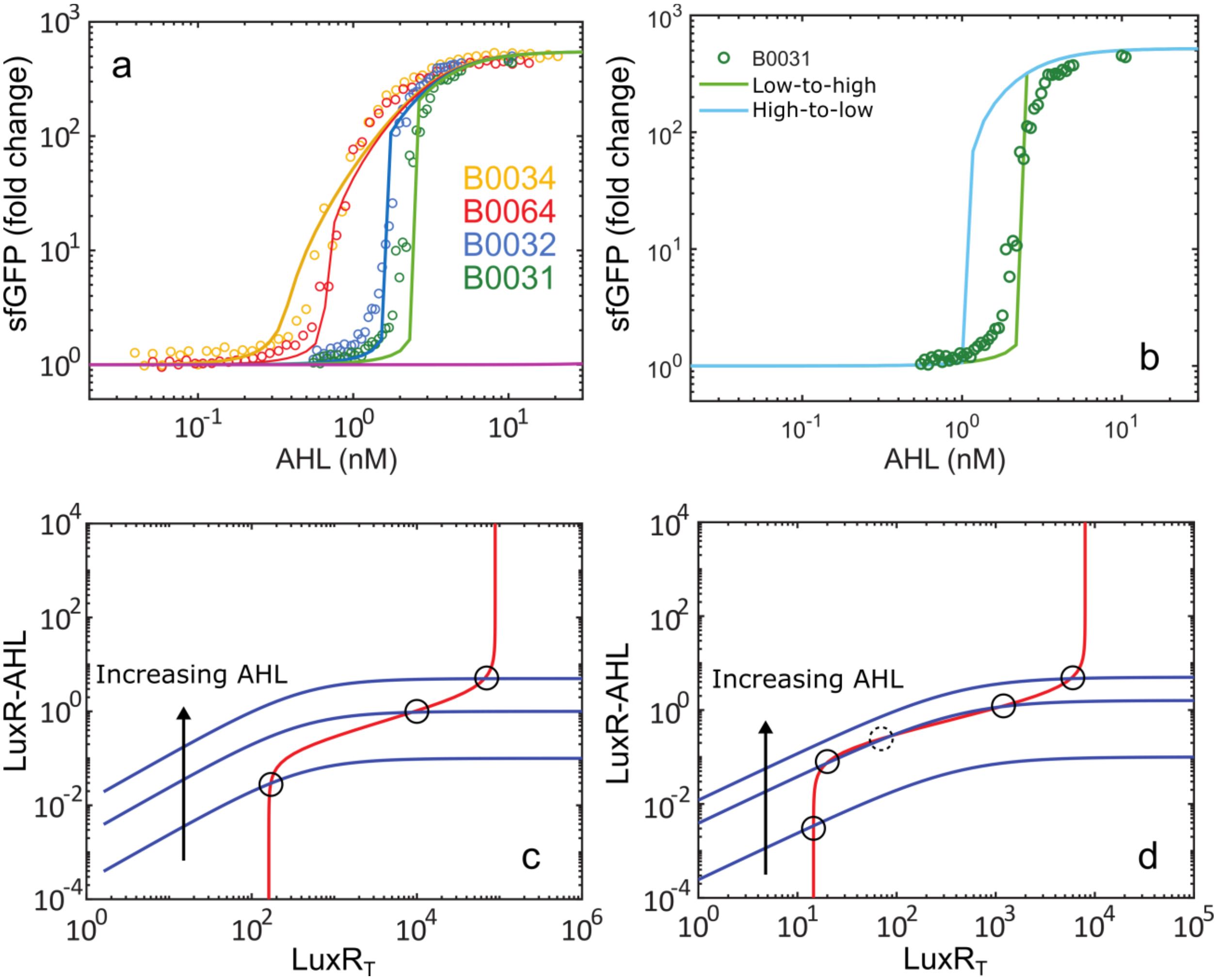

**Supplementary Figure 2. A deterministic kinetic model describes LuxR-based positive feedback IPs.** (**a**) Model fits (lines) to experimental mean sfGFP data (hollow points) as a function of AHL and RBS strength (colors) in LuxR-based positive feedback IPs. With a very low *luxR* translation rate (magenta), there is no activation of the IP. (**b**) Solutions for the system steady-state predict hysteresis in the AHL-mean transfer function for weak RBSs (B0031). (**c,d**) Visualization of the model at steady-state separated into two modules: (i) a transcription module (red line) shows total LuxR ($LuxR_T$) as a function of LuxR-AHL, and (ii) an interaction module (blue lines) shows LuxR-AHL as a function of $LuxR_T$ for a given concentration of AHL. Intersections (circles) are the system's steady-state. (**c**) For a strong RBS (yellow curve in (**a**)), there is always one stable steady-state, and the response increases smoothly as AHL increases. (**d**) For a weak RBS (green curve in (**a**)), at intermediate AHL induction, the system has two stable (solid-line circles) and one unstable (dashed-line circle) steady states (middle blue curve), making transitions extremely sharp and noisy.

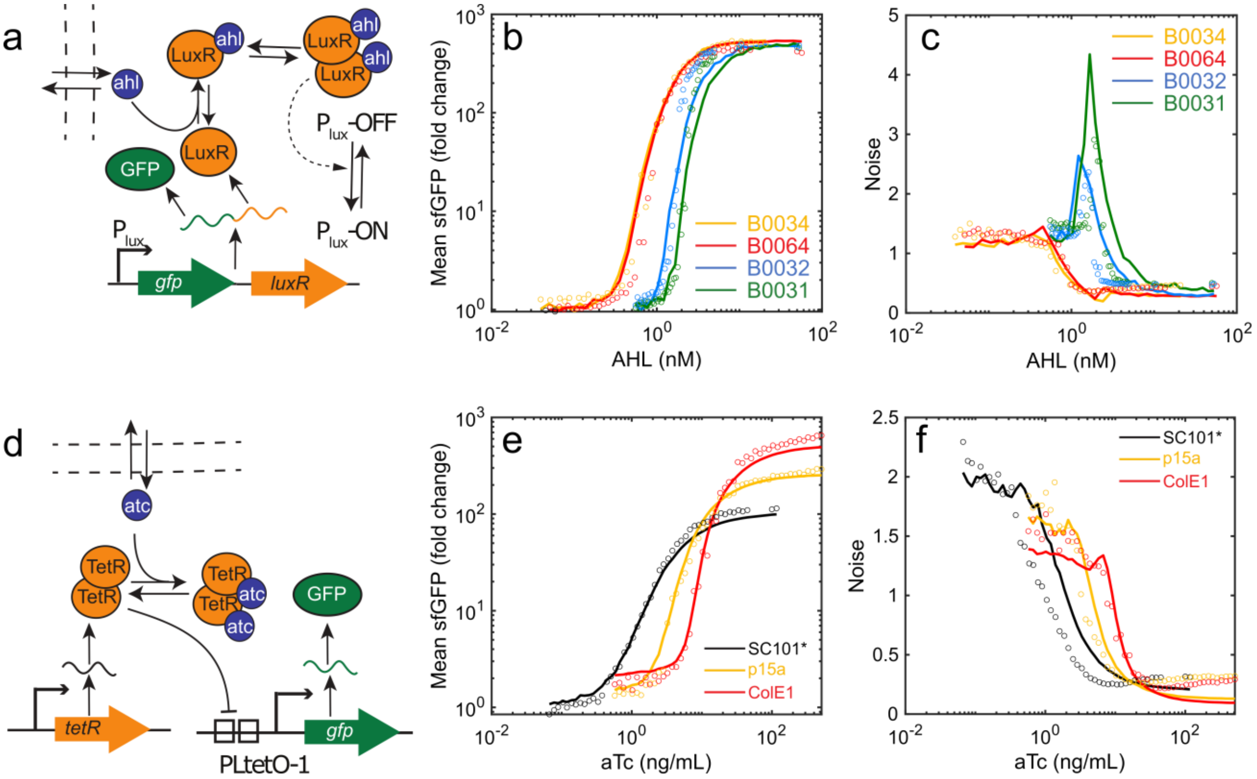

**Supplementary Figure 3. Stochastic kinetic models capture the behavior of low and high noise IPs in E. coli.** (**a**) Schematic representation of the model used to describe the LuxR based IPs for high noise sfGFP expression. AHL diffuses across the cell membrane, and extracellular AHL concentration is assumed constant. AHL binds LuxR monomers which then form dimers. AHL-LuxR dimers modulate the $P_{lux}$ promoter between ON and OFF states. In the ON state, LuxR and sfGFP genes are transcribed, resulting in transcriptional bursts and translation of sfGFP and LuxR from a single bicistronic mRNA. LuxR translation rate varies with RBS strength, resulting in different mean and noise characteristics. (**b**,**c**) Experimental (filled circles) and simulated (solid lines) sfGFP mean (**b**) and coefficient of variation (**c**) as a function of AHL and LuxR RBS. (**d**) Schematic representation of the model used to describe the TetR based IPs for low noise sfGFP expression. aTc diffuses across the cell membrane, and extracellular aTc concentration is assumed constant. Two aTc molecules bind TetR dimers sequentially. Free TetR dimers bind two $P_{LtetO-1}$ operator sites independently. The three promoter states (unbound, singly occupied, and doubly occupied) have varying transcription rates. TetR and $P_{LtetO-1}$ copy numbers are assumed to be 10, 25, and 50 for SC101*, p15a, and ColE1 based plasmids. (**e**,**f**) Experimental (circles) and simulated (solid lines) mean (**e**) and coefficient of variation (**f**) of sfGFP as a function of extracellular aTc concentration and plasmid copy number.

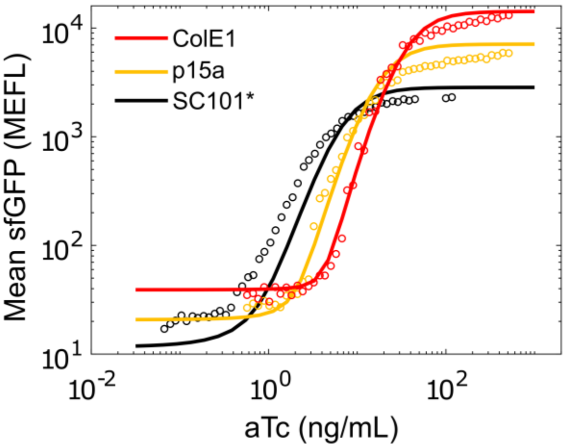

**Supplementary Figure 4. A deterministic kinetic model describes TetR based IP behavior.** aTc-mean transfer functions for TetR based no-feedback IPs with different origins of replication. Experimental values (circles) are overlaid with deterministic kinetic model predictions (smooth lines).

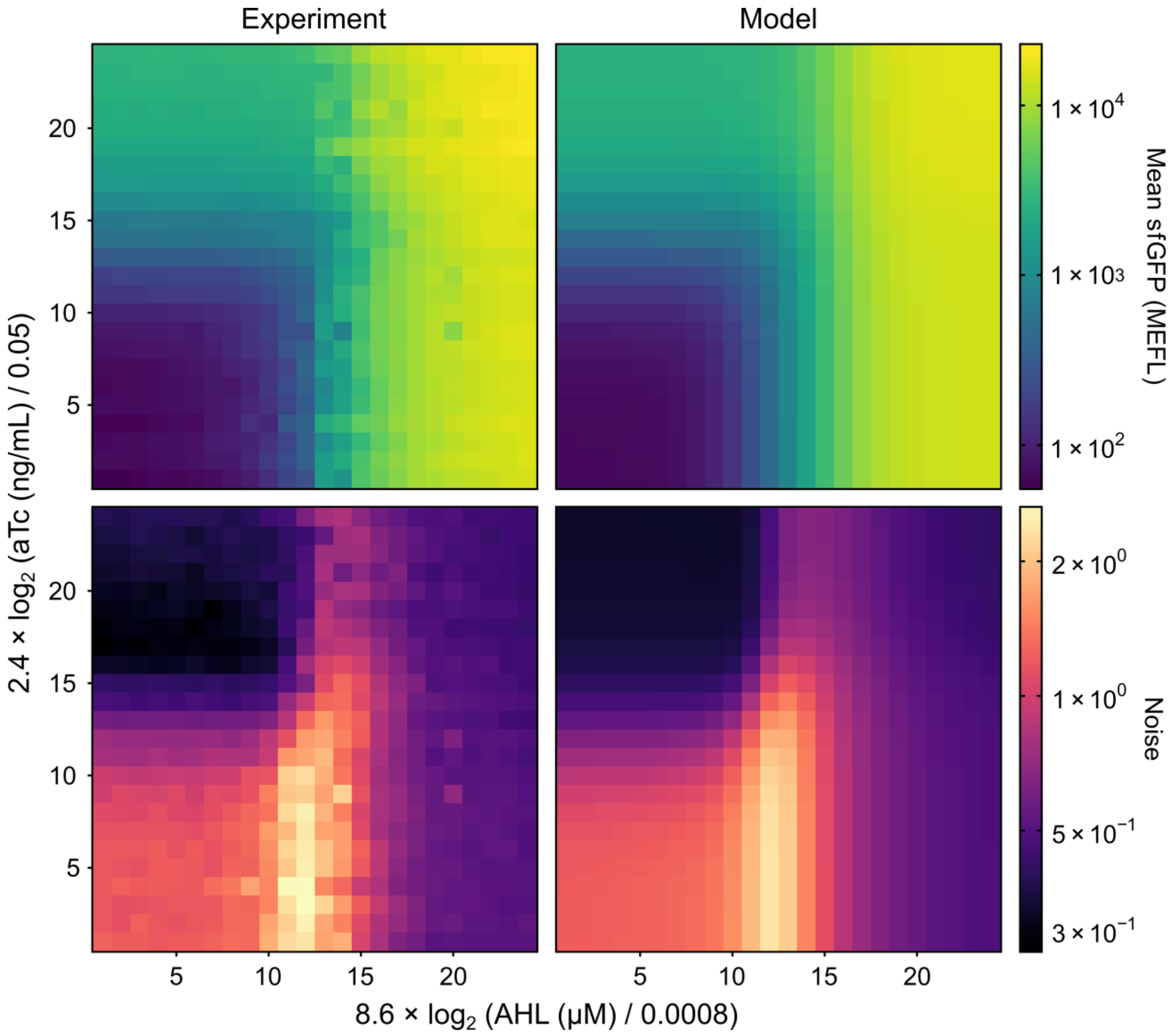

**Supplementary Figure 5. Characterization and modeling of $IP_h$/$IP_l$ mean and noise.** Heatmap visualization of experimentally measured (left panels) and model-predicted (right panels) steady-state mean (top panels) and noise (bottom panels) of cell populations co-expressing sfGFP from $IP_l$ and $IP_h$. Cells were exposed to exponentially distributed combinations AHL and aTc. Data with 0 AHL or aTc not shown due to log-scale. “Experiment” tiles are single replicates collected over three separate experiments performed on three separate days.

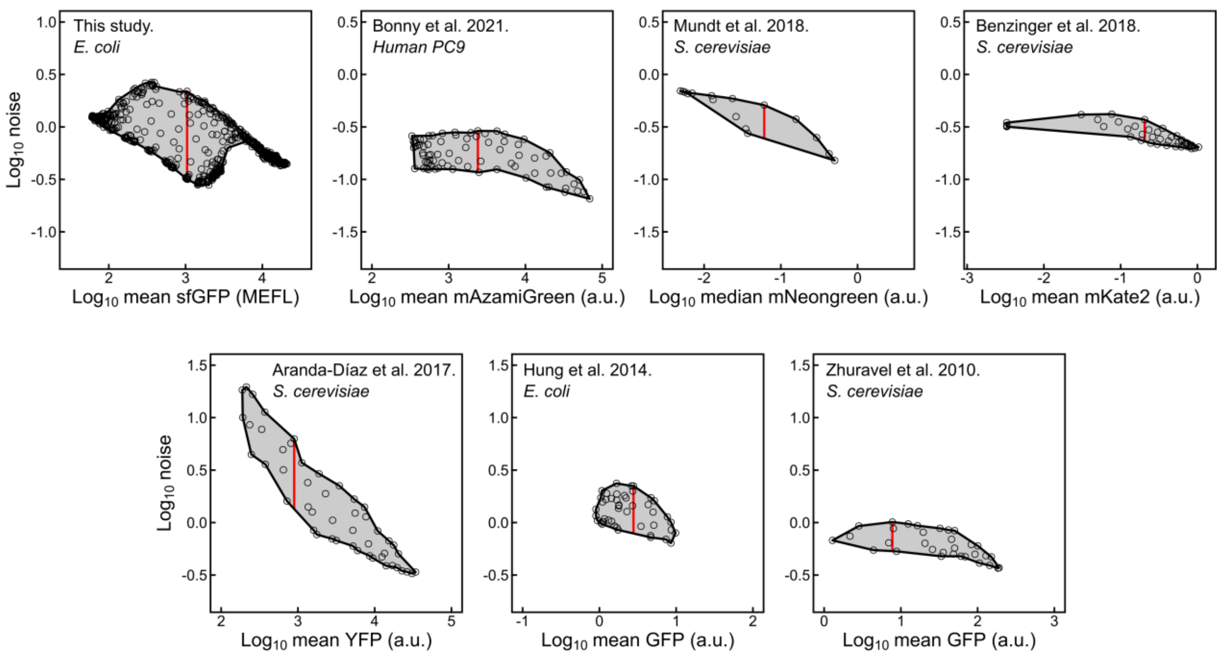

**Supplementary Figure 6. Visual comparison of mean-noise control systems from this and previous studies.** Log$_{10}$-space concave hulls (black lines), $\log_{10} F_A$ (shaded areas), and $\log_{10} F_\eta$ (red vertical cords) of point sets (circles) from this and previously described mean-noise control systems. Point sets of previous studies were digitally extracted from figures in their respective publications. All plots show 3-orders of magnitude in the x-dimension and 2.25-orders of magnitude in the y-dimension.

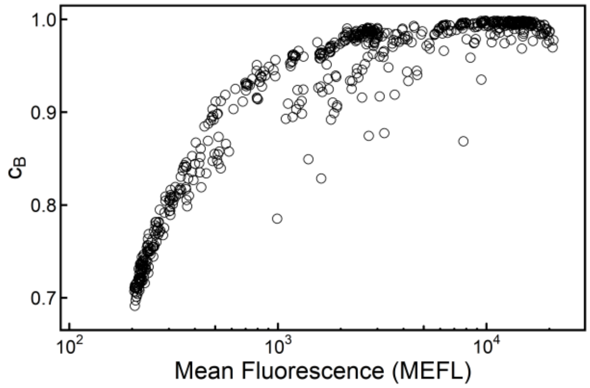

**Supplementary Figure 7. Bhattacharyya coefficient dependence on mean fluorescence.** Bhattacharyya coefficients of experimental-simulated fluorescence distribution pairs plotted against the mean of the corresponding experimental fluorescence distribution. Bhattacharyya coefficient dependence on the mean reflects the effect of autofluorescence and fluorescence from leaky sfGFP expression being overestimated and systematically biasing the simulated populations.

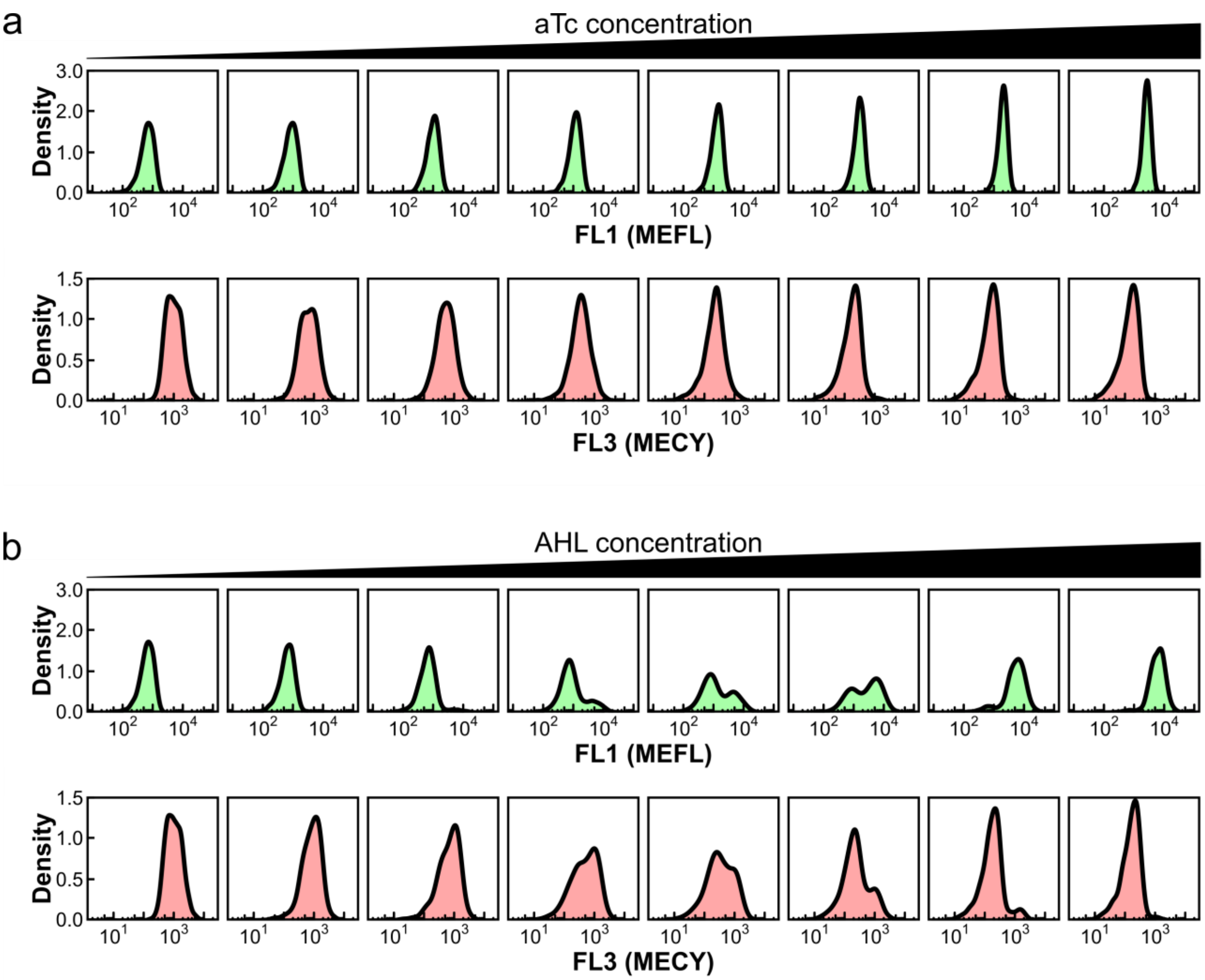

**Supplementary Figure 8. Comparison of sfGFP-PhlF$^{AM}$ and P$_{PhlF}$ output distributions.** FL1 and FL3 fluorescence distributions of cells transformed with pKG331-PhlF/pKG347-31-phlF/pKG361 and induced with varying amounts of aTc (**a**) or AHL (**b**). aTc concentrations: 0.00, 0.50, 0.67, 0.90, 1.20, 1.60, 2.14, 2.85 ng/uL; or AHL concentrations: 0.00, 0.043 0.051 0.060 0.071 0.084 0.192 0.315 µM were used to induce cultures.

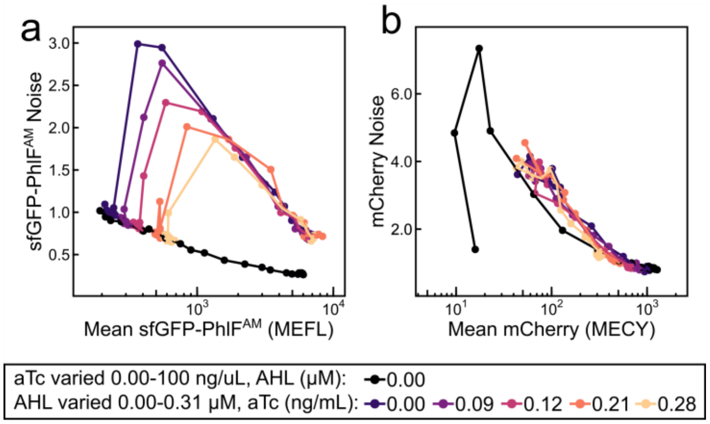

**Supplementary Figure 9. Noise in PhlF$^{AM}$ shows little evidence of propagation to $P_{PhlF}$.** Steady-state mean-noise transfer functions for (**a**) sfGFP-PhlF$^{AM}$ and (**b**) mCherry with exposure to combinations of AHL and aTc. Compared to sfGFP-PhlF$^{AM}$, mCherry noise is higher overall and shows only modest differences at the same mean between different induction conditions. Behavior that suggests most noise transmitted from PhlF$^{AM}$ to mCherry is filtered out or represents only a small portion of the total mCherry noise.

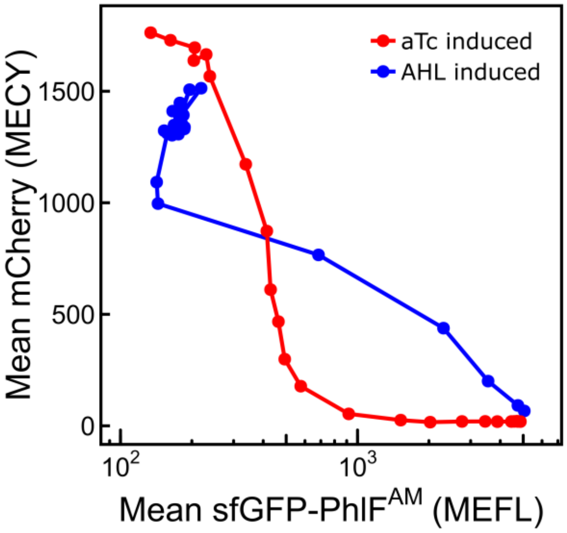

**Supplementary Figure 10. Shape of transcriptional output depends on transcription factor noise.** Mean mCherry expressed under the $P_{\mathit{PhlF}}$ promoter as a function of mean sfGFP-PhlF$^{AM}$ expression induced from $IP_l$ (red) or $IP_h$ (blue). MG1655 cells transformed with pKG331-PhlF/pKG347-31-phlF/pKG361 were grown under exposure to varying amounts of AHL or aTc.

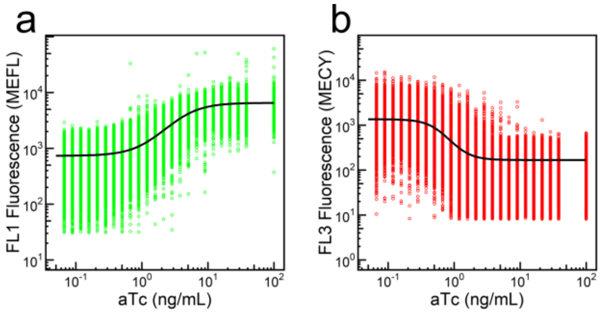

**Supplementary Figure 11. Single-cell inducer transfer functions for sfGFP-PhlF$^{AM}$ and $P_{PhlF}$-mCherry.** Fits to single-cell FL1 (**a**) and FL3 (**b**) fluorescence measurements of cell populations carrying pKG331-SC101*-phlF/pKG347-31-phlF/pKG361 when exposed to different amounts of aTc. Fits from (**a**) and (**b**) are used to construct a single cell transfer function between FL1 and FL3.

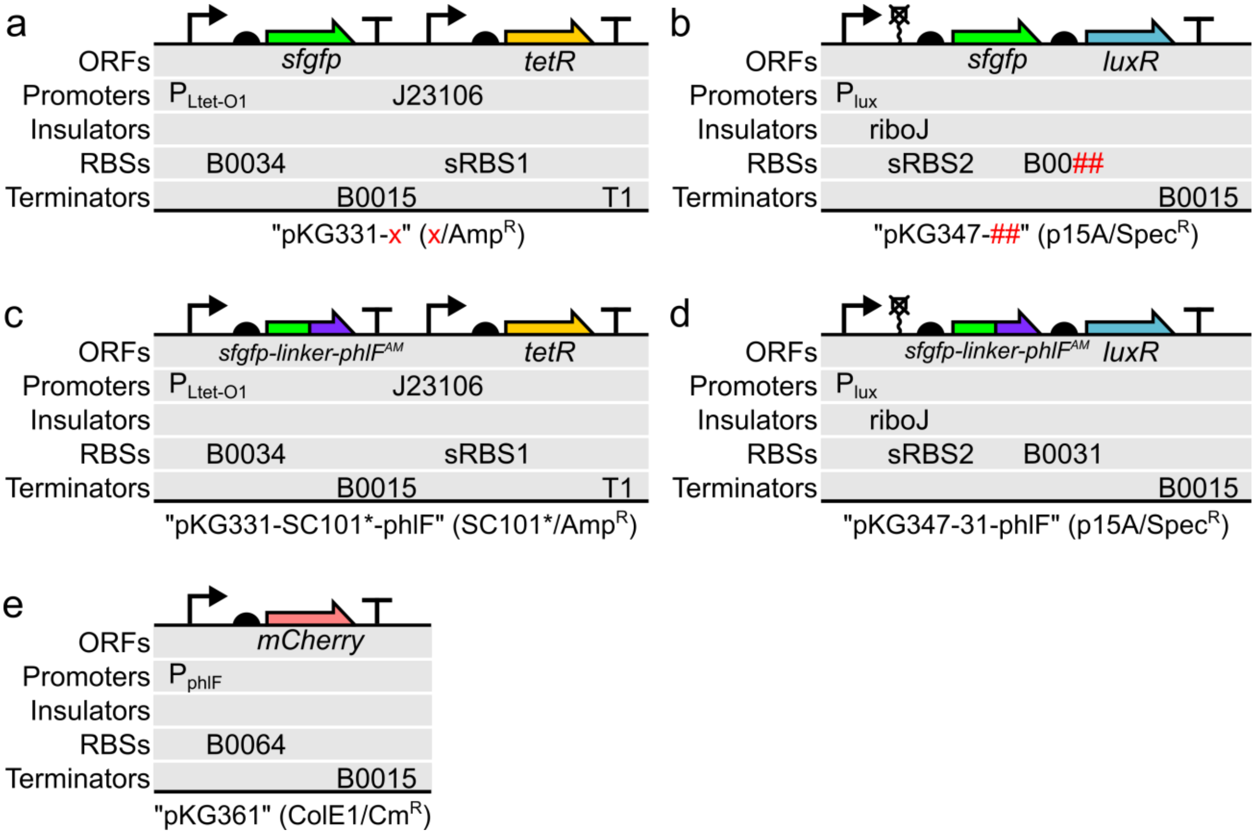

**Supplementary Figure 12. Plasmid maps.** (**a**) Low noise, TetR based plasmids used in this study. Origin of replication variants were used where “x” = ColE1, p15A, or SC101*. sRBS1 is a synthetic RBS designed using the RBS calculator with a predicted translation rate of 19027 a.u. (**b**) High noise, LuxR based plasmids used in this study. RBS variants were used where ## = 34, 64, 32, 31, or 33 (strongest to weakest). sRBS2 is a synthetic RBS designed using the RBS calculator with a predicted translation rate of 342 a.u. SC101* (**c**) and B0031 (**d**) variants of (**a**) and (**b**) with addition of a six amino acid linker (GGGGGH) and *phlF*$^{AM}$ to the C-terminus of the *sfgfp* open reading frame. (**e**) $P_{phlF}$ reporter plasmid.

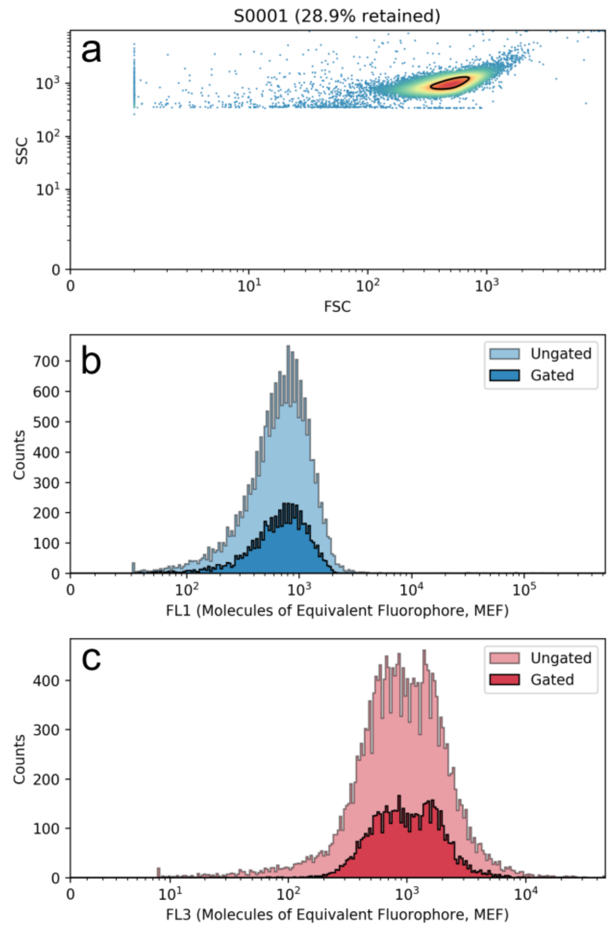

**Supplementary Figure 13. Density gating of flow cytometry data.** (**a**) Forward scatter (FSC) and side scatter (SSC) of a representative *E. coli* sample. Each point represents a particle (cell, cell aggregate, or debris) detected by the flow cytometer. Color indicates density (number of particles with a similar range of FSC and SSC). The black line represents the density gate and surrounds the highest density area containing 30% of the particles detected by the flow cytometer. An SSC threshold is applied below a level characteristic of the bacterial population such that events below the threshold (debris) are not detected. The first 250 events, final 100 events, and events with values in the first or last bin of the FSC, SSC, FL1, or FL3 channels were removed. (**b** and **c**) Cell fluorescence histograms of all particles in the forward/side scatter plot above, before and after applying the density gate.

### Supplementary Tables

| | LuxR RBS | RBS Strength | $a$ (MEFL) | $n_H$ | $k$ (μM) | $b$ (MEFL) |
|---|---|---|---|---|---|---|
| Best Fit Values | B0034 | 1 | 20480 | 4.27 | 0.001687 | 51.4 |
| | B0064 | 0.35 | 18410 | 5.22 | 0.001553 | 51.1 |
| | B0032 | 0.30 | 14920 | 10.00 | 0.002315 | 48.4 |
| | B0031 | 0.07 | 14310 | 10.40 | 0.002973 | 45.5 |
| | B0033 | 0.01 | 8560 | 4.54 | 0.725 | 31.3 |
| Standard Error | B0034 | 1 | 940 | 0.17 | 0.000066 | 2.2 |
| | B0064 | 0.35 | 780 | 0.20 | 0.000050 | 2.0 |
| | B0032 | 0.30 | 720 | 0.44 | 0.000042 | 2.2 |
| | B0031 | 0.07 | 890 | 0.50 | 0.000059 | 2.0 |
| | B0033 | 0.01 | 730 | 0.41 | 0.059 | 4.2 |

**Supplementary Table 1. Best fit parameters and standard errors for LuxR-based IP transfer functions.** RBS strength values are based on those reported by the Registry of Standard Biological Parts.

| Parameter | Description | Unit | Value |
|---|---|---|---|
| $LuxR_0$ | Basal expression of LuxR | a.u. | Varied 0.2-1.2 |
| $f$ | Maximal fold change per promoter copy | - | 458 |
| $K$ | Half activation concentration of LuxR-AHL | a.u. | 2.325 |
| $K_a$ | LuxR-AHL dissociation constant | a.u. | 7.5 |

**Supplementary Table 2. Parameter values for the deterministic kinetic model of LuxR based IPs.**

| Parameter | Description | Units | Value |
|---|---|---|---|
| $k_{dif}$ | Diffusion of AHL across membrane | $s^{-1}$ | 0.6 |
| $k_{dis}$ | $[LuxR\text{-}AHL]$ dissociation rate | $s^{-1}$ | $1.6\times10^{-3}$ |
| $K_a$ | $[LuxR\text{-}AHL]$ dissociation constant | copies/cell | 0.5 |
| $k_{on}^{0}$ | Switching ON rate for two-state promoter | $s^{-1}$ | $1\times10^{-3}$ |
| $k_{off}$ | Switching OFF rate for two-state promoter | $s^{-1}$ | 1.2 |
| $K$ | Dissociation constant for $[LuxR\text{-}AHL]$ to promoter | copies/cell | 300 |
| $f$ | Maximum fold increase in switching ON rate for two-state promoter | - | 916 |
| $k_{tr}$ | Transcription rate in the ON state | mRNA molecules $s^{-1}$ | 0.013 |
| $k_{tlnG}$ | Translation rate of GFP | $s^{-1}$ | 0.39 |
| $k_{tlnR}$ | Translation rate of LuxR | $s^{-1}$ | Ranges from (35-250)*$k_{md}$ |
| $K_{md}$ | mRNA decay rate | $s^{-1}$ | $4\times10^{-3}$ |
| $k_{pd}$ | Dilution rate due to growth | $s^{-1}$ | $3.1\times10^{-4}$ |
| $N$ | Mean promoter copy number $(p_{on} + p_{off})$ | molecules $cell^{-1}$ | 20 |
| $k_{in}[AHL(nM)]_{out}$ | Influx of AHL from the medium | molecules $s^{-1}$ | $0.04[AHL(nM)]_{out}$ |

**Supplementary Table 3. Parameter values for the stochastic kinetic model of LuxR based IPs.**

| Deterministic model | Stochastic model | Description |
|---|---|---|
| $LuxR_0$ | $\frac{N\frac{k_{tr}k_{tlnR}}{k_{pd}k_{md}}\left(\frac{k_{on}^0}{k_{on}^0+k_{off}}\right)}{\frac{k_{in}}{k_{pd}}}$ | Basal expression of LuxR (AHL = 0) |
| $K$ | $\approx \frac{K}{\sqrt{2}\frac{k_{in}}{k_{pd}}}$ | Threshold LuxR-AHL concentration for half-maximal activation of promoter |
| $K_a$ | $\frac{K_a}{\frac{k_{dif}}{k_{pd}}}$ | LuxR-AHL dissociation constant |
| $f$ | $\approx \frac{f}{2}$ | Maximal fold activation of LuxR promoter |

**Supplementary Table 4. Corresponding parameters between deterministic and stochastic kinetic models of LuxR based IPs.**

| | Origin of replication | $a$ (MEFL) | $n_H$ | $k$ (ng/mL) | $b$ (MEFL) |
|---|---|---|---|---|---|
| Best Fit Values | SC101* | 2088 | 2.046 | 4.74 | 18.87 |
| | p15a | 4640 | 2.84 | 14.02 | 24.4 |
| | ColE1 | 10220 | 3.36 | 25.9 | 34.9 |
| Standard Error | SC101* | 46 | 0.037 | 0.15 | 0.45 |
| | p15a | 210 | 0.13 | 0.82 | 2.0 |
| | ColE1 | 430 | 0.12 | 1.1 | 1.6 |

**Supplementary Table 5. Best fit parameters and standard errors for TetR based IP transfer functions.**

| Parameter | Description | Units | Value |
|---|---|---|---|
| $\alpha$ | Basal expression of GFP per promoter copy | a.u. | 0.76 |
| $f$ | Maximal fold change in GFP per promoter copy | | 370 |
| $N$ | Average promoter copy number | a.u. | [10, 25, 50] |
| $K_{tet}$ | Half-repression concentration of TetR | a.u. | 0.028 |
| $K_a$ | aTc-TetR dissociation constant | a.u. | 0.266 |
| $\beta$ | Constitutive expression of TetR per promoter copy | a.u. | 0.0751 |

**Supplementary Table 6. Parameters values for the deterministic kinetic model of TetR based IPs.**

| Parameter | Description | Units | Value |
|---|---|---|---|
| $k_{tettr}$ | Transcription of TetR | $s^{-1}$ | $1×10^{-4}$ |
| $k_{tln}$ | Translation of mTetR to TetR dimers | $s^{-1}$ | $6×10^{-2}$ |
| $k_{b1}$ | First aTc molecule binding $TetR_2$ | $molecule^{-1}$ $s^{-1}$ | $2.2×10^{-4}$ |
| $k_{d1}$ | aTc-$TetR_2$ dissociation | $s^{-1}$ | $5×10^{-6}$ |
| $k_{b2}$ | Second aTc molecule binding $TetR_2$ | $molecule^{-1}$ $s^{-1}$ | $2.2×10^{-4}$ |
| $k_{d2}$ | $aTc_2$-$TetR_2$ dissociation | $s^{-1}$ | $5×10^{-6}$ |
| $k_{f1}$ | TetR dimer binding $P_{LtetO-1}$ site 1 | $molecule^{-1}$ $s^{-1}$ | $5.8×10^{-4}$ |
| $k_{r1}$ | TetR dimer dissociating from site 1 | $s^{-1}$ | $2×10^{-5}$ |
| $k_{f2}$ | TetR dimer binding $P_{LtetO-1}$ site 2 | $molecule^{-1}s^{-1}$ | $5.8×10^{-4}$ |
| $k_{r2}$ | TetR dimer dissociating from site 2 | $s^{-1}$ | $2×10^{-5}$ |
| $k_0$ | Rate of transcription from $p_0$ | $s^{-1}$ | $6.8×10^{-4}$ |
| $k_1$ | Rate of transcription from $p_1$ | $s^{-1}$ | $2.2×10^{-6}$ |
| $k_2$ | Rate of transcription from $p_2$ | $s^{-1}$ | $1.4×10^{-6}$ |
| $k_{tlnG}$ | GFP translation rate | $s^{-1}$ | $9×10^{-2}$ |
| $k_{dif}$ | aTc diffusion rate across the membrane | $s^{-1}$ | $6.2×10^{-4}$ |
| $k_{pd}$ | Protein dilution due to growth | $s^{-1}$ | $3.1×10^{-4}$ |
| $k_{md}$ | mRNA decay rate | $s^{-1}$ | $2×10^{-3}$ |
| $k_{in}$ | Influx of aTc into cells | molecule $s^{-1}$ | $0.028[aTc_{ext}]$ |

**Supplementary Table 7. Parameters values for the stochastic kinetic model of TetR based IPs.**

| Deterministic | Stochastic | Description |
|---|---|---|
| $K_a = \frac{k_d + k_{dil}}{k_b}$ | $\sim \frac{k_{d1} + k_{pd}}{k_{b1}}$ | aTc-TetR dissociation constant |
| $\frac{TetR}{K_{tet}}$ | $\sim \sqrt{\frac{TetR_2}{K_{D1}}}$ | Half-repression concentration of TetR. $K_{D1} = \frac{\left(k_{r1} + k_{pd}\right)}{k_{f1}}$ |
| $[TetR]_T = \beta N$ | $[TetR_2]_T = N \frac{k_{tettr} k_{tlnR}}{k_{pd} k_{md}} + p_1 + 2p_2$ | Total TetR concentration |
| $f$ | $\frac{k_0}{\frac{k_1 + k_2}{2}}$ | Fold expression change between fully repressed and unrepressed states |

**Supplementary Table 8. Corresponding parameters between the deterministic and stochastic kinetic models of TetR based IPs.**

| Parameter ratios | Deterministic | Stochastic |
|---|---|---|
| $TetR_T\ (aTc = 0, N = 10)/K_{tet}$ | 27 | 14 |
| $K_a/K_{tet}$ | 9.5 | 1.86 |
| $f$ | 370 | 377 |

**Supplementary Table 9. Comparison of parameter values between the deterministic and stochastic kinetic models of TetR based IPs.**

| | Parameter | Best Fit Value | Standard Error |
|---|---|---|---|
| Low noise IP | $b$ | 10 | 10 |
| | $a$ | 2564 | 44 |
| | $n_H$ | 2.053 | 0.037 |
| | $k$ | 6.38 | 0.16 |
| | $c_0$ | 0.0925 | 0.0049 |
| | $c_1$ | 61.5 | 6.7 |
| | $c_2$ | 0 (Fixed) | - |
| | $c_3$ | 0 (Fixed) | - |
| | $c_4$ | 0 (Fixed) | - |
| High noise IP | $b$ | 60 | 10 |
| | $a$ | 13740 | 160 |
| | $n_H$ | 9.74 | 0.10 |
| | $k$ | 0.003016 | 0.000011 |
| | $c_0$ | 0.1974 | 0.0079 |
| | $c_1$ | 89 | 15 |
| | $c_2$ | 5.57 | 0.36 |
| | $c_3$ | 0.002137 | 0.000015 |
| | $c_4$ | 0.000258 | 0.000010 |

**Supplementary Table 10. Best fit parameters and standard errors for $IP_l/IP_h$ phenomenological mean and noise transfer functions.**

| Reference | $F_A$ | $F_\eta$ | $\alpha$ |
|---|---|---|---|
| This study | 11.39 | 6.88 | 0.212 |
| Bonny et al. 2021[1] | 5.29 | 2.47 | 0.5 |
| Mundt et al. 2018[2]* | 2.48* | 2.08* | 1.4 |
| Benzinger et al. 2018[3] | 2.22 | 1.61 | 4.0 |
| Aranda-Diaz et al. 2017[4] | 10.17 | 4.66 | 0.3 |
| Hung et al. 2014[5] | 2.29 | 2.84 | 0.3 |
| Zhuravel et al. 2010[6] | 2.76 | 1.89 | 0.9 |

**Supplementary Table 11. Performance metrics of mean-noise control systems.** Mean-noise pointsets from previous studies were digitally extracted from figures and analyzed as described in the main text. Alpha parameter ($\alpha$) was chosen manually.

*This dataset was reported in terms of median rather than mean and excluded extrinsic noise by normalizing fluorescence to a constitutive reporter.

| Parameter | Best Fit Value | Standard Error |
|---|---|---|
| $a$ | 5785 | 10 |
| $n_H$ | 1.8035 | 0.0098 |
| $k$ | 4.101 | 0.014 |
| $b$ | 734.1 | 4.8 |
| $c$ | -1187.9 | 4.6 |
| $m_H$ | 2.763 | 0.043 |
| $j$ | 0.5786 | 0.0037 |
| $d$ | 1355.5 | 3.7 |

**Supplementary Table 12. Best fit parameters and standard errors for single-cell inducer transfer functions for sfGFP-PhlF$^{AM}$ and P$_{phlF}$-mCherry.**

| Plasmid | Primary Features | Resistance |
|---|---|---|
| pKG331-ColE1 | TetR, no feedback, ColE1 | Amp$^{R}$ |
| pKG331-p15a | TetR, no feedback, p15a | Amp$^{R}$ |
| pKG331-SC101* | TetR, no feedback, SC101* | Amp$^{R}$ |
| pKG347-34 | LuxR, + feedback, B0034 | Spec$^{R}$ |
| pKG347-64 | LuxR, + feedback, B0064 | Spec$^{R}$ |
| pKG347-32 | LuxR, + feedback, B0032 | Spec$^{R}$ |
| pKG347-31 | LuxR, + feedback, B0031 | Spec$^{R}$ |
| pKG347-33 | LuxR, + feedback, B0033 | Spec$^{R}$ |
| pKG331-SC101*-phlF | TetR, no feedback, ColE1 | Amp$^{R}$ |
| pKG347-31-phlF | LuxR, + feedback, B0031 | Spec$^{R}$ |
| pKG361 | $P_{phlF}$ with mCherry output | Cm$^{R}$ |

**Supplementary Table 13. Names of plasmids used in this study.**

### Supplementary Methods

### Deterministic kinetic modeling of TetR and LuxR based IPs

Models for both TetR and LuxR IPs share a common core: an inducer molecule binds a transcription factor (TF). The TF dimerizes either in the inducer bound state (LuxR) or in the free state (TetR) and binds promoter elements and activates (LuxR) or represses (TetR) transcription of *gfp* (Supplementary Figure 3). The gene for the TF itself is either under the control of the same regulated promoter (LuxR) or expressed constitutively (TetR). With this common core of reactions, using approximations, we construct relatively simple deterministic models that capture the mean GFP dose-response (normalized to basal state) over variations in TF synthesis rate (LuxR IPs) or total promoter copy number (TetR IPs). The simplicity allows us to explore analytical solutions of the TetR IP kinetic model that helps explain how the steepness of mean GFP dose-response changes with increasing mean plasmid copy number. The analytical solution for the kinetic model of the LuxR IP allows us to uncover the mechanism behind a steep, step-like transfer function of GFP when using weak RBSs.

*Deterministic kinetic model of TetR based IPs*

We simplify the above-described core of reactions for the TetR IPs with the following assumptions: GFP is expressed from the promoter with basal/leaky expression independent of TetR-repression (with rate $\alpha$ per promoter copy per generation) and an additional unrepressed level that is $f$-fold higher when TetR is not bound to the promoter. Therefore, GFP expression rate per promoter copy depends on the concentration of free TetR available to repress, so that:

$$\frac{d[GFP]}{dt} = \alpha k_{dil} N \left( 1 + \frac{f}{1 + \left(\frac{[TetR]}{K_{tet}}\right)^2} \right) - k_{dil}[GFP] \tag{7}$$

Here, $N$ denotes the mean plasmid copy number and $K_{tet}$, the concentration of TetR at which the promoter is repressed with 50% probability, denotes the affinity of TetR molecules to the promoter. $k_{dil}$ denotes a growth/dilution rate as we assumed GFP (and other proteins) to be stable.

The dynamics of TetR can be described similarly. TetR is expressed constitutively at a constant level of $\beta$ per promoter copy per generation and diluted by cell growth. In addition, posttranslational regulation reactions include TetR binding (rate constant $k_b$) and dissociation (rate constant $k_d$) of aTc:

$$\frac{d[TetR]}{dt} = \beta k_{dil} N - k_b [aTc][TetR] + k_d [aTc\text{-}TetR] - k_{dil}[TetR] \quad (8)$$

$$\frac{d[aTc\text{-}TetR]}{dt} = k_b [aTc][TetR] - k_d [aTc\text{-}TetR] - k_{dil}[aTc\text{-}TetR] \quad (9)$$

In steady state, we can use Eq. (9) to express the concentration of the aTc-TetR complex as:

$$[aTc\text{-}TetR] = \frac{[aTc][TetR]}{K_a} \quad (10)$$

where $K_a = \frac{k_d + k_{dil}}{k_b}$. Furthermore, we assume fixed total aTc and TetR concentrations:

$$[aTc]_T = [aTc] + [aTc\text{-}TetR] \quad (11)$$

$$[TetR]_T = [TetR] + [aTc\text{-}TetR] \quad (12)$$

In steady state, we can find the total concentration of TetR by summing Eqs. (8) and (9) to obtain $[TetR]_T = N\beta$. From 9 & 11, we solve for free TetR as a function of the total concentration of TetR and free aTc:

$$[TetR] = \frac{[TetR]_T}{1 + \frac{[aTc]}{K_a}} \quad (13)$$

Substitution in Eq. (11) results in a quadratic that can be solved to express concentration of free aTc as a function of total concentrations of TetR and aTc:

$$\frac{[aTc]}{K_a} = \frac{1}{2}\left(-\left(1+\frac{[TetR]_T}{K_{tet}}-\frac{[aTc]_T}{K_a}\right)+\sqrt{\left(1+\frac{[TetR]_T}{K_{tet}}-\frac{[aTc]_T}{K_a}\right)^2+\frac{4[aTc]_T}{K_a}}\right) \quad (14)$$

Thus, for a given plasmid copy number and inducer concentration, we can compute steady-state concentration of free TetR from Eqs. (13) and (14). Using this concentration, we find steady-state expression of GFP given by setting $\frac{d[GFP]}{dt} = 0$:

$$GFP = \alpha N\left(1+\frac{f}{1+\left(\frac{[TetR]}{K_{tet}}\right)^2}\right) \quad (15)$$

Solving Eqs. (13) and (14) for each value of $N$ = {10, 25, 50} for plasmids {SC101*, p15a, ColE1}, over a range of inducer concentrations gives us steady-state mean GFP transfer function. Parameters for this model {$K_{tet}$, $f$, $\alpha$, $\beta$, $K_a$} were fit to the normalized experimental data for the three plasmid variants using the function `particleswarmoptimization` in MATLAB (Supplementary Table 6). We normalize GFP fluorescence to the value at 0 aTc for the smallest copy number and fit the above parameters with arbitrary units.

To understand how steepness of the transfer function depends on the plasmid copy-number, we can analytically obtain an expression for the slope of dose-response at half repression ($[TetR]/K_{tet} = 1$) from Eqs. (13) – (15):

$$\left(\frac{\partial[GFP]}{\partial[aTc]_T}\right)_{[TetR]=K_{tet}} = \left(\frac{\partial[GFP]}{\partial[TetR]}\frac{\partial[TetR]}{\partial[aTc]}\right)\left(\frac{\partial[aTc]}{\partial[aTc]_T}\right) = \left(\frac{\alpha f K_{tet}}{2K_a}\right)\left(\frac{\frac{\beta N}{K_{tet}}}{\frac{\beta N}{K_{tet}}+\frac{K_{tet}}{K_a}}\right) \quad (16)$$

The above is a monotonically increasing function of $N$, saturating at high values. Thus, as promoter copy numbers increase, the transfer function of the IP becomes steeper, consistent with experimental results (Supplementary Figure 4).

*Deterministic kinetic model of LuxR based IPs*

For the LuxR based IPs, GFP, LuxR, and LuxR-AHL concentrations can be modeled similarly to the TetR based IPs (Eqs. (7) – (9)). However, since both LuxR and GFP are transcribed from the same promoter, we don't track GFP concentration dynamics explicitly but instead assume GFP concentration is proportional to the total concentration of LuxR. LuxR is synthesized from its mRNA through LuxR-AHL binding the promoter (positive autoregulation) and dilutes due to growth in cell volume. No active degradation of LuxR is assumed. In contrast to the TetR IP, we do not vary the plasmid copy numbers in the LuxR based IPs. Therefore, we do not explicitly include a variable for the promoter copy number in LuxR equations. Instead, we combine parameters for promoter copy number, transcription rate, translation rate, mRNA degradation rate, and protein dilution into a single parameter, $LuxR_0$, representing basal concentration of LuxR in the absence of AHL. This parameter is varied in the deterministic model to understand the effect of changes in the RBS strength. The dynamics of LuxR and LuxR-AHL can be described by the following equations:

$$\frac{d[LuxR]}{dt} = k_{dil}LuxR_0 - k_b[AHL][LuxR] + k_d[LuxR\text{-}AHL] - k_{dil}[LuxR] \tag{17}$$

$$\frac{d[LuxR\text{-}AHL]}{dt} = k_b[AHL][LuxR] - k_d[LuxR\text{-}AHL] - k_{dil}[LuxR\text{-}AHL] \tag{18}$$

At steady state, therefore:

$$[LuxR\text{-}AHL] = \frac{[LuxR][AHL]}{K_a} \tag{19}$$

where $K_a = \frac{k_d + k_{dil}}{k_b}$, and:

$$[LuxR]_T = LuxR_0 \left( \frac{1 + f\left(\frac{[LuxR\text{-}AHL]}{K}\right)^2}{1 + \left(\frac{[LuxR\text{-}AHL]}{K}\right)^2} \right) \tag{20}$$

Conservation relations for LuxR (neglecting LuxR-AHL molecules bound to the promoter) and AHL gives:

$$[LuxR]_T = [LuxR] + [LuxR\text{-}AHL] \tag{21}$$

$$[AHL]_T = [AHL] + [LuxR\text{-}AHL] \tag{22}$$

As with Eq. (14), solving these three equations for LuxR-AHL:

$$[LuxR\text{-}AHL] = \frac{1}{2}\left((K_a + [AHL]_T + [LuxR]_T) - \sqrt{(K_a + [AHL]_T + [LuxR]_T)^2 - 4[AHL]_T[LuxR]_T}\right) \tag{23}$$

The steady-state concentration of total LuxR for a given inducer concentration lies at the intersection of Eqs. (20) and (23). Thus, we can compute steady-state total LuxR concentration as a function of total AHL amount by solving Eqs. (20) and (23) simultaneously. This model is described by four parameters (Supplementary Table 4): $LuxR_0$, $f$, $K$, $K_a$.

*Condition for ultrasensitivity to AHL concentration*

The sensitivity of total LuxR concentration (proportional to GFP) to inducer concentration can be defined as a dimensionless parameter characterizing the slope in logarithmic space, i.e., $S = \frac{d\log[LuxR]_T}{d\log[AHL]_T}$.

If Eq. (20) is represented as $[LuxR]_T = F([LuxR\text{-}AHL])$, and Eq. (23) as $[LuxR\text{-}AHL] = G([AHL]_T, [LuxR]_T)$, then S can be obtained using the chain rule of differentiation:

$$S = \frac{L(F, [LuxR\text{-}AHL])L(G, [AHL]_T)}{1 - L(F, [LuxR\text{-}AHL])L(G, [LuxR]_T)} \tag{24}$$

Where $L(a, b) = \frac{\partial \log a}{\partial \log b}$ represents log gain of $a$ with respect to $b$. Ultrasensitivity then exists when the product of log gains $L(F, [LuxR\text{-}AHL])L(G, [LuxR]_T) \rightarrow 1$. Graphically, this condition can be visualized in Supplementary Figure 2. When the denominator in Eq. (24) is negative, the steady state becomes unstable. This corresponds to bistability – two stable steady states separated by an unstable one, i.e., three intersection points. This results in a sharp increase in LuxR and GFP as inducer concentration increases (Supplementary Figure 2b) as the system traverses from the low steady state to the high. In contrast (at high $LuxR_0$, i.e., with a strong RBS), either $L(F, [LuxR\text{-}AHL])$ or $L(G, [LuxR]_T)$ is $\ll 1$ at any inducer concentration (Supplementary Fig. 2c), resulting in a monostable system with no ultrasensitivity.

### Stochastic kinetic modeling of TetR and LuxR based IPs

The common core of reactions was expanded to elementary reactions to perform simulations using Gillespie SSA. Elementary reactions of the TetR based IPs were based on a previously published model[7] in *S. cerevisiae* as a starting point.

#### *Stochastic kinetic model of TetR based IPs*

We assume aTc diffuses into cells with constant flux proportional to aTc concentration in the medium, and aTc diffuses out of cells with first-order kinetics. We assume TetR translates directly into a dimer (dimerization is fast and stable) and aTc molecules bind each TetR dimer with identical rates. We assume TetR dimers bind each operator site independently, and operator sites are knocked free every cell doubling during plasmid replication. This leads us to the following set of reactions:

$$\phi \xrightarrow{k_{in}} aTc$$

$$aTc \xrightarrow{k_{dif}} \phi$$

$$p_{js} \xrightarrow{k_{tettr}} p_{js} + mTetR$$

$$mTetR \xrightarrow{k_{tln}} TetR_2 + mTetR$$

$$aTc + TetR_2 \xrightarrow{k_{b1}} aTc\text{-}TetR_2$$

$$aTc\text{-}TetR_2 \xrightarrow{k_{d1}} aTc + TetR_2$$

$$aTc + aTc\text{-}TetR_2 \xrightarrow{k_{b2}} aTc_2\text{-}TetR_2$$

$$aTc_2\text{-}TetR_2 \xrightarrow{k_{d2}} aTc + aTc\text{-}TetR_2$$

$$TetR_2 \xrightarrow{k_{dil}} \phi$$

$$aTc\text{-}TetR_2 \xrightarrow{k_{dil}} \phi$$

$$aTc_2\text{-}TetR_2 \xrightarrow{k_{dil}} \phi$$

$$p_0 + TetR_2 \xrightarrow{k_{f1}} p_1$$

$$p_1 \xrightarrow{k_{r1}} p_0 + TetR_2$$

$$p_1 + TetR_2 \xrightarrow{k_{f2}} p_2$$

$$p_2 \xrightarrow{k_{r2}} p_1 + TetR_2$$

$$p_2 \xrightarrow{k_{dil}} p_0 + 2\,TetR_2$$

$$p_1 \xrightarrow{k_{dil}} p_0 + TetR_2$$

$$p_0 \xrightarrow{k_0} p_0 + mGFP$$

$$p_1 \xrightarrow{k_1} p_1 + mGFP$$

$$p_2 \xrightarrow{k_2} p_2 + mGFP$$

$$mGFP \xrightarrow{k_{md}} \phi$$

$$mTetR \xrightarrow{k_{md}} \phi$$

$$mGFP \xrightarrow{k_{tlnG}} GFP + mGFP$$

$$GFP \xrightarrow{k_{dil}} \phi$$

We set the initial promoter copy number for both GFP ($p_0$) and TetR ($p_{js}$) to one of {10, 25, 50}. Next, we set a value for the influx rate of aTc ($k_{in}$) as input. We run simulations using the Gillespie SSA algorithm in the SimBiology package in MATLAB. The length of the simulation was chosen $>> 1/k_{dil}$ (~40 generations) to ensure steady state is reached. The end state of GFP from 500 such simulations was used to obtain mean and variance statistics.

*Stochastic kinetic model of LuxR based IPs*

We constructed a stochastic model of the LuxR based IPs by expanding the core reactions from the deterministic model in the same way as the TetR stochastic model. A stochastic model for an AHL-LuxR based system has been described in the literature for *Vibrio harveyi* quorum-sensing and was used as a reference[8]. We assumed elementary reactions for LuxR-AHL dimers binding promoter copies and upregulating transcription rate of *luxR* and *gfp*. This model was able to match the noise characteristics of all four RBS strengths without extrinsic noise. However, we could not match the transfer function's steepness for the two weak RBSs (B0031 and B0032). We could rescue this fit by increasing the basal transcription rate from the promoter, but this resulted in a much lower CV at 0 AHL for any RBS, as expected for a Poisson process[9,10]. We believe this to result from bistability at low RBS strengths and the associated critical slowdown[11]. Given that our purpose for the stochastic models was to confirm that the model can explain noise properties over varying RBS strengths, we opted not to study this model further. Instead, we opt for a two-state promoter[10] to account for the burstiness of transcription.

Rather than assuming transcription rate at two levels: basal and active (similar to the TetR IPs above), we consider a two-state promoter that undergoes stochastic switches between OFF and ON states. Transcription occurs in bursts at a single rate in the ON state, while the OFF state is non-permissible to transcription[12,13]. It has been shown that transcription time-series properties are largely independent of the particular regulation scheme of a promoter[12]. Therefore, we do not explicitly model elementary reactions for transcription factors binding. Instead, we assume the propensity of the promoter to switch from OFF to ON states (i.e., the frequency of bursts) is a saturating function of the transcription factor concentration as shown in Eq. (25). It is worth noting that the model functions equally well if $k_{off}$ (i.e. duration of bursts) is assumed to be a decreasing function of LuxR-AHL.

$$k_{on} = k_{on}^{0}\left(\frac{1+f\left(\frac{[LuxR\text{-}AHL]}{K}\right)^{2}}{1+\left(\frac{[LuxR\text{-}AHL]}{K}\right)^{2}}\right) \tag{25}$$

We assume a constant influx of AHL proportional to $AHL_{out}$, whereas outflux is proportional to intracellular free AHL concentration. With these assumptions, we arrive at the following set of reactions:

$$\phi \xrightarrow{k_{in}[AHL(nM)]_{out}} AHL$$

$$AHL \xrightarrow{k_{dif}} \phi$$

$$AHL + LuxR \xrightarrow{k_b} LuxR\text{-}AHL$$

$$LuxR\text{-}AHL \xrightarrow{k_{dis}} AHL + LuxR$$

$$p_{OFF} \xrightarrow{k_{on}} p_{ON}$$

$$p_{ON} \xrightarrow{k_{off}} p_{OFF}$$

$$p_{ON} \xrightarrow{k_{tr}} p_{ON} + m$$

$$m \xrightarrow{k_{tlnG}} GFP$$

$$m \xrightarrow{k_{tlnR}} LuxR$$

$$m \xrightarrow{k_{md}} \phi$$

$$GFP \xrightarrow{k_{pd}} \phi$$

$$LuxR \xrightarrow{k_{pd}} \phi$$

$$LuxR\text{-}AHL \xrightarrow{k_{pd}} \phi$$

An exact deterministic analytical solution can be derived for this stochastic model such that it is consistent with the deterministic model (Eqs. (29) and (30)).

To account for extrinsic noise, we allow variability in protein decay ($k_{pd}$), mRNA decay ($k_{md}$), transcription rate ($k_{tr}$) and promoter copy number ($N$). For each run, we select a value for the above four parameters from a log-normal distribution around mean values (using CV of 0.15) listed in Supplementary Table 3. With no AHL in the system, we simulate a pre-AHL steady state of ~10hrs. For this, we run the SSA 200 times in MATLAB. The mean of each species is rounded off and considered as the initial condition for the next step. Next, at each AHL concentration, we fix the flux of AHL diffusing into the cell as $k_{in}[AHL(nM)]_{out}$. We then simulate 500 runs for 6 h starting from the pre-AHL initial condition. The end copy numbers of GFP are used to compute mean and variances to make the plots in Supplementary Fig. 4 B and C.

### Corresponding parameters between the stochastic and deterministic models

*TetR based IPs*

A few differences exist between the deterministic and stochastic model formulations that need to be accounted for while comparing parameters. While the deterministic model assumes concentration of aTc in the medium (i.e. input aTc concentration) is equal to intracellular concentration, the stochastic model assumes a constant influx of aTc into cells. Furthermore, TetR is assumed to have a strong dimerization dissociation constant such that nearly all copies exist as dimers. We then consider two independent operator sites for TetR dimers to bind to the promoter[14]. To understand which parameter combinations of the stochastic model correspond to the four parameters of the deterministic model (GFP synthesis rate does not need to be compared as we fit models to normalized GFP curves), we can evaluate approximate deterministic steady-state expressions of all the species.

The procedure followed to arrive at parameter combinations of the stochastic model that correspond to the deterministic model parameters is the same as derived in detail in the next section for the LuxR models. In brief, we describe the reactions used for stochastic simulations as ODEs and solve at steady state. We obtain three equations corresponding to Eqs. (13) – (15) in the deterministic TetR model. Where direct comparison is not possible or analytical equations are cumbersome, we make approximations using the parameter values used for stochastic simulations (Supplementary Table 7). The parameter comparisons are listed in Supplementary Tables 8 and 9.

Since a TetR dimer ($TetR_2$) with a single aTc molecule bound is incapable of repressing the promoter, we compare $K_a$ values directly (the effective dissociation constant of aTc and TetR). To find a comparable half-repression concentration of TetR ($K_{tet}$), we can compute analytical expressions for the three states ($p_0$, $p_1$, and $p_2$) of the promoter as a function of free TetR dimers, and then write the expression for GFP expression as a function of $p_0$, $p_1$, and $p_2$ to compare with Eq. 15. However, this equation is not amenable for direct comparison. Here, we make use of our parameter values (in this case, TetR dissociation from promoter sites is assumed much slower than dilution due to growth, i.e. $k_{r1} \ll k_{pd}$) to reduce the expression such that it allows for a direct comparison to Eq (15). To find the parameter corresponding to $f$ in the deterministic model, we can assume $p_1$ (singly repressed) and $p_2$ (fully repressed) states of the promoter express at a single averaged rate (i.e., if $k_1/k_2$ = 1.5 fold, then we assume $p_1 + p_2$ expresses transcripts at $1.25 \times k_2$, while $p_0$ expresses at $k_0$).

To fully specify the kinetic parameters for the stochastic model, we assume protein degradation/dilution rate ($k_{pd}$) on the order of cell doubling time (~35 min), mRNA half-life ($k_{md}$) of ~5 min, and GFP and TetR translation rates ($k_{tlnG}$ and $k_{tln}$) of 0.09 and 0.06

(i.e. mean translation burst sizes of 45 and 30) respectively[15–17]. Transcription rate of TetR mRNA was obtained by fit and lies in a biologically feasible range. The transcription rates of GFP mRNA from $p_0$, $p_1$ and $p_2$ states of the promoter are constrained by the coefficient of variation of GFP observed at 0 aTc. The ratios between $p_0$, $p_1$ and $p_2$ state transcription rates are also constrained by the observed maximum GFP fold-change. Having specified translation, mRNA and protein degradation rates, and a mean plasmid copy number of ~10 for SC101*, we can estimate initial transcription rate values such that a CV of ~2 can be expected from the stochastic simulations[18–20]. Thus, we specify rate parameters for the transcription and translation processes modeled. Dissociation rate of aTc-TetR has been reported to be much slower than dilution due to growth[21] and we assumed it to be ~1/100 of $k_{pd}$ (consistent with a model of aTc-TetR in S. cerevisiae[7]). The forward rates of aTc binding free and singly bound TetR dimers were assumed to be equal and were calculated based on reported dissociation constant ranges for the aTc-TetR complex[23]. Association and dissociation rate constants for TetR dimers binding either binding site of $P_{Ltet\text{-}O1}$ were assumed equal and independent of each other. Dissociation rate on the order of $10^{-5}$ $s^{-1}$ has been reported between TetR and tetO2 sites[23,24], and the bimolecular forward rate of association is constrained by fit and is within the range for a typical cell volume of 1 $um^3$. Finally, diffusion of tetracyclines across cell membrane occurs at ~1 h timescales[25].

*LuxR based IPs*

One major difference between the stochastic and deterministic models is in accounting for AHL. The deterministic model assumes the total intracellular concentration of AHL to be the same as the input (external AHL). Thus, in order to find the equation corresponding to Eq. (23) i.e. LuxR-AHL as a function of external AHL and total LuxR concentration, we first need to relate steady state intracellular concentration of AHL to the external concentration. Therefore, we represent the set of reactions used for stochastic simulations by ordinary differential equations describing the concentrations of AHL, LuxR, and LuxR-AHL. From these equations, we can write the rate of change of total intracellular concentration of AHL as

$$\frac{d[AHL]}{dt} + \frac{d[LuxR\text{-}AHL]}{dt} = k_{in}[AHL]_{out} - k_{dif}[AHL] - k_{pd}[LuxR\text{-}AHL] \qquad (26)$$

We can substitute $[LuxR\text{-}AHL] = \frac{[LuxR][AHL]}{K_a}$ at steady state, where $K_a = \frac{k_{dis}+k_{pd}}{k_b}$. Therefore, at steady state

$$\frac{k_{in}}{k_{pd}}[AHL]_{out} = [AHL]\left(\frac{k_{dif}}{k_{pd}} + \frac{[LuxR]}{K_a}\right) \qquad (27)$$

This relates intra- and extra-cellular AHL concentrations. From this point, the same steps used for obtaining solutions to the deterministic model of LuxR based IPs can give us an analytical solution for this more detailed model. The conservation relation for LuxR gives us:

$$[LuxR]_T = [LuxR]\left(1 + \frac{[AHL]}{K_a}\right) \tag{28}$$

Substituting the expression for AHL above and solving for LuxR-AHL gives us Eq. (29) corresponding to Eq. (23) for the deterministic model. Note that here we use the notation $[AHL]_T$ to denote $[AHL]_{out}$ and not the total intracellular AHL concentration.

$$\frac{[LuxR\text{-}AHL]}{\phi_2} = \frac{1}{2}\left(\left(\frac{\phi_1 K_a}{\phi_2} + [AHL]_T + \frac{[LuxR]_T}{\phi_2}\right) - \sqrt{\left(\frac{\phi_1 K_a}{\phi_2} + [AHL]_T + \frac{[LuxR]_T}{\phi_2}\right)^2 - 4[AHL]_T\frac{[LuxR]_T}{\phi_2}}\right) \tag{29}$$

Where $K_a = \frac{k_{dis}+k_{pd}}{k_b}$, $\phi_1 = \frac{k_{dif}}{k_{pd}}$, $\phi_2 = \frac{k_{in}}{k_{pd}}$. In order to enable a direct comparison to Eq. (23), we have scaled all other parameters except $[AHL]_T$ in the stochastic model. Thus, $K_a$ in Eq. (23) corresponds to $\frac{\phi_1 K_a}{\phi_2}$ in the stochastic model.

To derive an expression for total concentration of LuxR (i.e. an equation corresponding to Eq. (20)), we note that for a constitutively active two state promoter, mean mRNA copy number is given by $< mRNA > = \frac{k_{TX}}{k_{md}}\left(\frac{k_{on}}{k_{on}+k_{off}}\right)$ [12], and the corresponding protein copy number by $\frac{k_{tln}}{k_{pd}} < mRNA >$. Where $k_{TX}$ the transcription rate in the ON state of the promoter. Substituting in $k_{TX}$ as $Nk_{tr}$, and the $[LuxR - AHL]$ dependence assumed in Eq. (25), we obtain:

$$[LuxR]_T = LuxR_0 \left( \frac{1 + f\left(\frac{[LuxR\text{-}AHL]}{K}\right)^2}{1 + \left(1 + \frac{f k_{on}^0}{k_{off}}\right)\left(\frac{[LuxR\text{-}AHL]}{K}\right)^2} \right) \quad (30)$$

Where $LuxR_0 = N\frac{k_{tr}k_{tlnR}}{k_{pd}k_{md}}\left(\frac{k_{on}^0}{k_{on}^0+k_{off}}\right)$. We note that $[LuxR - AHL]$ and $[LuxR]_T$ values need to be scaled by $\frac{1}{\phi_2}$ in Eq. (26) to enable direct comparison with the deterministic model. In order to obtain parameter combinations in Eq. (30) that correspond to the deterministic solution (Eq. (20)), we note that the maximum value of $[LuxR]_T$ in the equation above is $\approx LuxR_0\frac{f}{2}$ when $[LuxR\text{-}AHL] \gg K$ and $\frac{fk_{on}^0}{k_{off}} \approx 1$ (Supplementary Table 4). Thus, $f$ in Eq. (20) is equivalent to $f/2$ in the stochastic model. Next, for a value of $\frac{[LuxR\text{-}AHL]}{K} = \frac{1}{\sqrt{2}}$, Eq. (30) reduces to $[LuxR]_T \approx LuxR_0\frac{f}{4}$, which is half of the maximum value of $[LuxR]_T$. Thus, $K$ in Eq. (20) is equivalent to $\frac{K}{\sqrt{2}\,\phi_2}$ in the stochastic model. Similarly, $LuxR_0$ in Eq. (20) is equivalent to $\frac{LuxR_0}{\phi_2}$ in the stochastic model.

To specify the full list of parameters for the stochastic model, we assume $k_{pd}$ is given by a doubling time of ~35 mins, and the mean mRNA lifetime is ~3 mins. Maximal transcription rate ($1.3 \times 10^{-2}\ s^{-1}$) and mean number of proteins translated per transcript, i.e. translational burst size, (35-250) lie in biologically feasible ranges[15–17]. The two-state promoter is specified by $k_{on}^0,\ k_{off}$ and $f$, which are constrained such that $k_{off} \gg k_{on}^0$ [12,13] and $f$ is consistent with the deterministic model. The LuxR-AHL dissociation rate is chosen to be $\gg k_{pd}$ [8], and diffusion of AHL across membrane is assumed to be on the order of a few seconds to minutes[8,26]. The LuxR based IP is constructed on a p15a plasmid, for which we assume a mean copy number of 20 [14]. In our model, major sources of noise are the ON-OFF switching of the promoter leading to bursts in transcription, as well as translation of long-lived LuxR protein from short-lived mRNA (i.e. $k_{md} \gg k_{pd}$)[27,28]. Corresponding parameters between deterministic and stochastic models are summarized in Supplementary Table 4, and the full list of parameters for the stochastic model in Supplementary Table 3.

### Phenomenological fit for mean and noise of $IP_h/IP_l$

Eqs. (1) and (3) were fit to experimental data using phenomenological expressions for mean and noise of individual IPs as a function of inducer concentration (Figure 3b-d, Supplementary Fig. 6, Supplementary Table 7). While a discrete approach to solving Eqs. (1) and (3) using two experimentally measured distributions is possible (Figure 4), this phenomenological approach enables continuous prediction of mean and noise as a function of inducer concentrations.

We describe the steady-state mean of an individual IP with a Hill function:

$$\mu(i) = \frac{a i^{n_H}}{i^{n_H} + k^{n_H}} + b \qquad (31)$$

where $i$ is inducer concentration, $b$ is the amount of basally produced protein, $a$ is the mean output range, $k$ is the inducer concentration at which $\mu(i)$ reaches 50% of its range, and $n_H$ is the Hill coefficient. For individual IP noise, we adopt the following equation following the approach from a previous study[29]:

$$\eta^2(i) = c_1 + \frac{c_2}{\mu(i)} + \frac{c_3}{\left(1 + \left(\frac{i - c_4}{c_5}\right)^2\right)} \qquad (32)$$

where the first (constant), second (inverse scaling with the mean), and third (peak-shaped function) terms aim to capture extrinsic, intrinsic, and transmitted sources of noise, respectively.

Best fit parameters were found by plugging Eqs. (31) and (32), for each IP, into Eqs. (1) and (3) and fitting to experimental data in Figure 3.

#### Single-cell transfer functions for sfGFP-PhlF$^{AM}$ and $P_{PhlF}$-mCherry

We relate single-cell mCherry concentrations to single-cell sfGFP-PhlF$^{AM}$ concentrations through aTc transfer functions. To do this, we use the single-cell fluorescence data for populations that were exposed only to aTc. We fit Hill functions to the single-cell aTc-FL1 and aTc-FL3 transfer functions (Supplementary Fig. 10, Supplementary Table 11):

$$FL1(aTc) = \frac{a[aTc]^{n_H}}{[aTc]^{n_H} + k^{n_H}} + b \tag{33}$$

$$FL3(aTc) = \frac{c[aTc]^{m_H}}{[aTc]^{n_H} + j^{m_H}} + d \tag{34}$$

We then use these relationships to express FL3 as a function of FL1. Because Eq. (33) is asymptotically bound at $b$ and $a + b$, and individual cells stochastically reside outside these boundaries, we create a piecewise function:

$$FL3(FL1) = \begin{cases} d, & FL1 < b \\ f(FL1), & b \leq FL1 \leq a + b \\ d + c, & FL1 > a + b \end{cases} \tag{35}$$

where $f(FL1)$ is obtained by expressing Eq. (34) in terms of FL1.
